# A blueprint for mutation-defined hallmark vulnerabilities across human cancers

**DOI:** 10.1101/2025.10.24.684315

**Authors:** Ran Xu, Rebecca San Gil, Xin Tracy Liu, Yanfei Qi, Derek Tran, Chun Xu, Justin J-L Wong, Lenka Munoz, Graham J. Mann, Yuchen Feng

## Abstract

Hallmark gene mutations shape cancer cell vulnerabilities and inform drug discovery^1–3^. A systematic map of hallmark gene mutation-defined cancer dependencies and therapeutic responses is essential to uncover novel targets and refine therapeutic strategies. Here, we present the first pan-cancer blueprint of hallmark vulnerabilities, systematically linking hallmark gene mutation markers to cancer cell dependencies and drug sensitivities across 22 cancer cohorts. We integrated multi-omics data from patient tumours with large-scale CRISPR-Cas9 screens and pharmacologic profiling of over a thousand cancer cell lines. Our analysis revealed the cancer type-specific nature of hallmark gene expression programs, uncovered previously unrecognised mutation-target gene dependencies, and highlighted metabolic programs as a dominant class of functional vulnerabilities. Notably, we identified oxidative phosphorylation (OXPHOS) addiction in *CDKN2A*-loss lung squamous cell carcinoma (LUSC) and experimentally validated this dependency. Our validation highlights the greater selectivity of *CDKN2A*-loss LUSC cells to metformin, an FDA-approved antidiabetic drug known for its OXPHOS inhibitory activity. Proteogenomic integration further prioritised targets overexpressed in mutant tumours, constituting therapeutic windows. Pharmacologic profiling identified both oncology and non-oncology agents with selective activity in mutation-defined subgroups, revealing opportunities for drug repurposing. Our machine learning framework, Comet-X, for the first time fully leveraged gene mutation combinations to predict these target dependencies and drug responses. The resulting pan-cancer mutation-dependency map provides a comprehensive resource of hallmark gene targets and candidate therapeutics, stratified by mutation markers, to pave the way for drug development, clinical trial design and discovery research.

## Main text

Precision oncology aims to overcome the limitations of conventional cancer type-based treatment strategies by stratifying tumours into biologically distinct subtypes defined by molecular signatures that can predict the susceptibility of cancers to specific therapeutics^4,5^. At the core of precision oncology is the concept that somatic mutations driving tumour initiation, progression and treatment resistance shape distinct cancer vulnerabilities that can be therapeutically targeted^6,7^. However, to date, most biomarker discovery efforts have focused on recurrent genomic alterations with matched targeted therapies or established chemotherapeutic regimens. For example, the PI3K inhibitor Inavolisib for *PIK3CA*-mutated advanced breast cancer and associations between *ADAMTS* mutations and improved chemotherapy sensitivity in high-grade serous ovarian carcinoma^8,9^. The broader landscape of mutation-associated vulnerabilities remains largely unexplored.

The hallmarks of cancer define core biological programs that drive tumorigenesis and progression, offering a conceptual compass to map cancer vulnerabilities^1^. Recent advances in large-scale multi-omics initiatives offer an unprecedented opportunity to define cancer vulnerabilities and mutation markers predictive of therapeutic responses^10,11^. Publicly available resources, including The Cancer Genome Atlas (TCGA), Clinical Proteomic Tumour Analysis Consortium (CPTAC), Cancer Cell Line Encyclopedia (CCLE), and the Cancer Dependency Map (DepMap), collectively provide extensive genomic, transcriptomic, proteomic, gene dependency, and pharmacologic sensitivity profiles from thousands of patient-derived tumours and cancer cell lines^10,12–14^. However, comprehensive and systematic characterization of hallmark gene mutation-associated functional dependencies and therapeutic sensitivities has yet to be established. Furthermore, current biomarkers for cancer vulnerabilities and therapeutic guidance are predominantly based on single-gene mutations^15^. As a result, vulnerabilities arising from combinatorial mutations, which are common across all cancer types, remain poorly characterized. Machine learning, capable of integrating complex, high-dimensional data, can be used to construct informative and predictive models for discovering combinatorial mutation-vulnerability associations^16,17^. Leveraging this computational tool could significantly expand our understanding of cancer vulnerabilities beyond single-gene analyses and uncover previously unrecognized therapeutic opportunities.

In this study, we integrated multi-omic and functional datasets to construct a comprehensive blueprint of hallmark gene mutation-associated dependencies and therapeutic sensitivities across 22 cancer types. Our analysis not only provides deeper insights into known cancer targets and therapeutics, but also systematically identifies and prioritizes novel, clinically actionable candidates, including opportunities for drug repurposing such as metformin for *CDKN2A*-mutant lung squamous cell carcinoma (LUSC) and digoxin for *SPTA1*-mutant stomach adenocarcinoma (STAD). The identified vulnerabilities span overexpressed hallmark proteins, essential functional programs and drug sensitivities linked to single-gene mutations. Most importantly, our innovative machine learning-based framework was able to consider mutation markers both singly and in unrestricted combinations to identify these targets and therapeutic opportunities. Collectively, this work offers a scalable resource of context-specific cancer vulnerabilities to inform future anti-cancer drug discovery and precision medicine strategies.

### Characterizing hallmark gene expression patterns across cancer types

To systematically map how hallmark gene mutations shape cancer cell vulnerabilities and therapeutic responses, we integrated large-scale multi-omics datasets spanning genomics, transcriptomics and proteomics, as well as gene dependency and drug response profiles across up to 30 cancer types (Fig. 1a). This included somatic mutation data, transcript and protein abundance, CRISPR-Cas9 loss-of-function screens, and oncology and non-oncology drug sensitivity profiles, sourced from TCGA (9,104 tumour and 732 normal samples), CPTAC (805 tumour and 523 normal samples), and Cancer Cell Line Encyclopedia (CCLE, 751 cancer cell lines) (Fig. 1a). By harmonizing these diverse datasets, we constructed a comprehensive blueprint of mutation-defined dependencies and pharmacologic sensitivities across diverse human cancers to uncover novel actionable targets and drug repurposing opportunities for precision oncology.

**Fig. 1.**
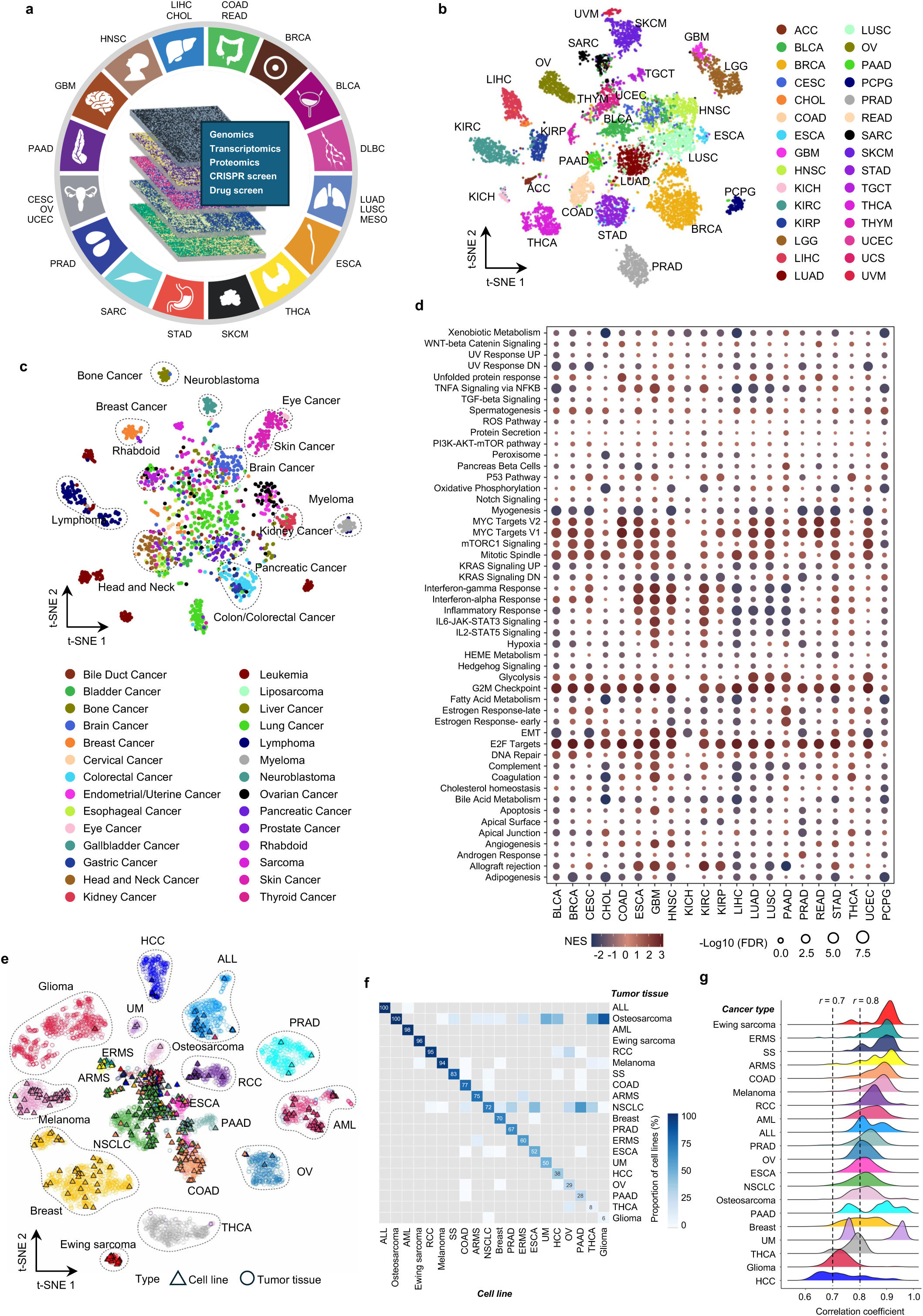

To investigate similarities and differences in hallmark gene expression patterns across cancer types, we analyzed RNA-seq data from TCGA patient-derived tumours and CCLE cell lines spanning 30 cancer types. The 4384 hallmark genes, representing 50 hallmark pathways, were obtained from the Human Molecular Signatures Database^18^. We applied *t*-distributed stochastic neighbor embedding (*t*-SNE) clustering to visualize the extent to which hallmark gene expression profiles vary across cancer types^19^. This analysis revealed lineage-specific clustering of both tumours and cell lines, with samples grouping according to their cancer type of origin (Fig. 1b, c). Consistently, gene set enrichment analysis (GSEA) of TCGA tumour versus matched normal transcriptomes also uncovered that each cancer type preferentially activated distinct hallmark pathways (Fig. 1d). For example, while MYC and E2F pathways were broadly activated across multiple cancers, UV response pathway showed selective enrichment in head and neck squamous cell carcinoma (HNSC) (Fig. 1d). Metabolic pathways, oxidative phosphorylation (OXPHOS) and glycolysis, were co-activated in LUSC, pancreatic adenocarcinoma (PAAD), and uterine corpus endometrial carcinoma (UCEC), whereas STAD and HNSC exhibited glycolysis enrichment alone (Fig. 1d)^20^. Together, these analyses uncover the cancer type-specific nature of hallmark gene expression programs, which lays the foundation for exploring their potential as lineage-specific therapeutic targets.

Given that investigations into hallmark gene dependencies rely on CRISPR-Cas9 screens in cancer cell lines, we first assessed whether cell lines serve as suitable models for inferring tumour-relevant dependencies. We integrated several large RNA-seq gene expression datasets including CCLE, TCGA, TARGET and Treehouse, and performed a joint dimensionality reduction analysis of cancer cell lines and patient-derived tumours using Celligner, an unsupervised alignment method, to evaluate the concordance of hallmark gene expression profiles between tumour and cell line cohorts^21^. This analysis revealed that transcriptional signatures of cell lines were largely aligned with tumour samples of the same cancer type with distinct cancer types forming separate clusters (Fig. 1e), recapitulating the lineage-specific patterns observed in tumour and cell line cohorts when analyzed individually (Fig. 1b, c). Notably, 15 out of 20 tested cancer types demonstrated high alignment, with over half of cell lines clustering with their tumour type of origin (Fig. 1e, f). The highest alignment was found for acute lymphoblastic leukemia (ALL, 100%), osteosarcoma (100%), and acute myeloid leukemia (AML, 98%), whereas thyroid carcinoma (THCA) and glioma exhibited the lowest alignment (<10%) (Fig. 1f, g). These findings confirm that hallmark gene expression profiles are broadly consistent between cancer cell lines and patient tumours but distinct across cancer types. This supports leveraging gene dependency data in cancer cell lines to systematically identify hallmark gene vulnerabilities and prioritize cancer type-specific therapeutic targets.

### Mapping mutation-associated gene dependencies

To profile hallmark gene dependencies in each cancer cohort, we analyzed genome-wide CRISPR-Cas9 screen data of 751 cancer cell lines representing 26 cancer types, of which three cell lines or more were available, obtained from the DepMap. This analysis uncovered hallmark genes whose knockout (KO) significantly impaired cell growth varied across diverse cancer types, ranging from 851 in UCS to 2,446 in HNSC (Fig. 2a, Extended Data Fig. 1a-c). Of these dependencies, 195 genes exhibited cancer type-specific essentiality, for example, KO of LDLRAP1 selectively reduced cell viability in lymphoid neoplasm Diffuse large B-cell lymphoma (DLBC) (Fig. 2b, Extended Data Fig. 1d), whereas 564 genes, such as *RAN*, were essential across all 26 cancer types (Fig. 2b, Extended Data Fig. 1e), representing pan-cancer vulnerabilities. Notably, we observed that cohort-level dependency calls did not fully capture intra-cohort heterogeneity, as some genes classified as essential showed minimal effects in a subset of cell lines (Fig. 2c), while others not identified as significant still exhibited strong effects in specific cell lines (Fig. 2d). These observations highlight the heterogeneous nature of gene dependencies within cancer types, underscoring the importance of incorporating molecular features, such as mutation markers, to stratify cancer cell lines and uncover context-specific vulnerabilities.

**Fig. 2.**
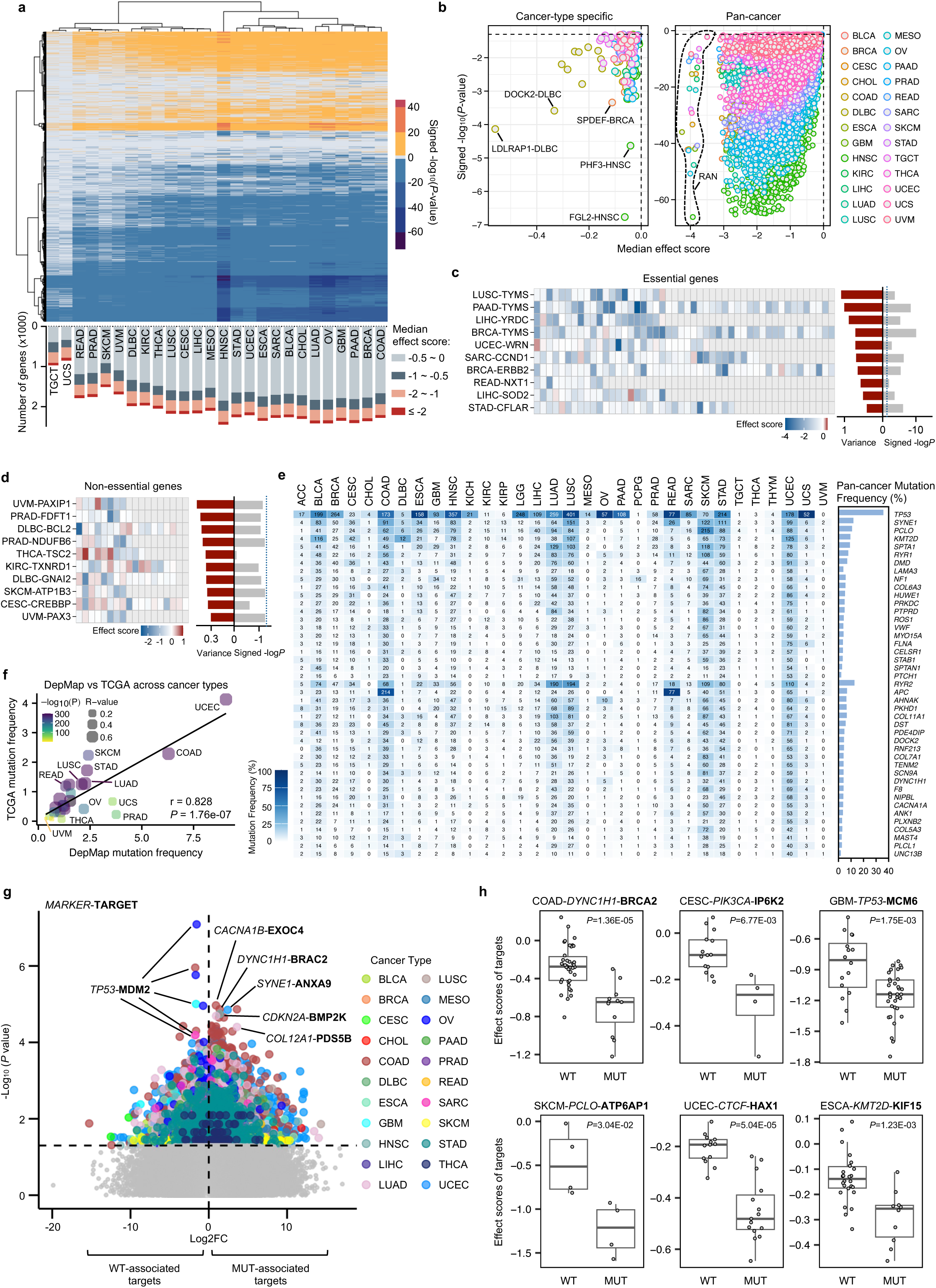

To integrate genetic context into dependency stratification, we next profiled hallmark gene mutation frequencies across cancer types using TCGA data (Extended Data Fig. 2a). Focusing on non-silent mutations, our analysis revealed distinct mutation patterns in different cancer types (Extended Data Fig. 2a). Across the pan-cancer cohort, *TP53* (35.8%), *RYR2* (12.8%) and *SYNE1* (11.9%) were the most frequently mutated hallmark genes (Fig. 2e, Extended Data Fig. 2b). However, mutation prevalence of these commonly altered genes varied substantially across cancer types. For example, although *TP53*, *SYNE1*, *PCLO*, *KMT2D*, *SPTA1*, and *RYR1* were commonly mutated in over 31 cancer types, their frequencies ranged from 0.2% to 91.2% (Fig. 2e, Extended Data Fig. 2c). In addition, some hallmark genes exhibited high mutation frequencies in specific cancer types, for instance, *APC* mutations were prevalent in colon adenocarcinoma (COAD, 73.8%) and rectum adenocarcinoma (READ, 85.6%), and *RYR2* mutations in lung adenocarcinoma (LUAD, 37%) and LUSC (40.4%) (Fig. 2e)^22,23^. Together, this analysis identified hallmark genes that were frequently mutated within each cancer cohort, serving as candidate molecular markers for context-specific dependency analysis. To assess the relevance of these mutations in cancer cell line models, we next examined hallmark gene mutation frequencies in DepMap cell lines (Extended Data Fig. 2d) and compared them with TCGA patient tumours. Mutation frequencies were highly correlated between tumours and cell lines in 25 out of 26 cancer types, with uveal melanoma (UVM) being the only exception showing weak concordance (Fig. 2f). Furthermore, evaluation of the overall mutation frequencies across the 26 cancer types revealed a strong correlation between TCGA and DepMap datasets (r = 0.828, *P* = 1.76e-07) (Fig. 2f), suggesting that hallmark gene mutation patterns were consistent between tumours and cell lines. Together with our previous finding that hallmark gene expression profiles were highly aligned between the two cohorts (Fig. 1e), these results reinforce the suitability of DepMap cell lines for modeling hallmark gene mutation-associated dependencies in downstream analyses.

To generate the marker-target dependency map, we defined markers as hallmark genes mutated in over 10% of cases within each cancer type, representing prevalent molecular features within the cohort (Extended Data Fig. 3a, b). This approach yielded 1-211 mutation markers across 24 cancer types, with UCEC having the most and kidney renal clear cell carcinoma (KIRC), thyroid carcinoma (THCA), and UVM having only one marker each (Extended Data Fig. 3a, b). We then integrated CRISPR-Cas9 screen and mutation data from DepMap to stratify cell lines into marker-mutant and marker-wildtype subgroups, which allowed systematic investigation of mutation-associated gene dependencies. This analysis uncovered 51 to 21,969 significant mutation marker-target associations across 22 cancer types, where KO of the target gene led to significantly greater inhibition of cell growth in marker-mutant cells compared to their wildtype counterparts (Fig. 2g, Extended Data Fig. 4a). Notably, among the top-ranked associations were MDM2 dependency in *TP53*-wildtype cells, consistent with its role as a negative regulator of the tumour suppressor p53 (Fig. 2g, Extended Data Fig. 4b), and PIK3CA dependency in *PIK3CA*-mutant cells, aligned with the known oncogenic function of *PIK3CA* mutations (Extended Data Fig. 4c)^24,25^. In addition to these well-characterized associations, our analysis also uncovered novel vulnerabilities associated with marker mutations (Fig. 2g, h). For example, in COAD, cells harboring *DYNC1H1* mutations exhibited increased sensitivity to KO of BRCA2, playing a central role for repairing DNA double-strand breaks by homologous recombination (Fig. 2h)^26^. In cervical squamous cell carcinoma and endocervical adenocarcinoma (CESC), knockout of inositol phosphokinase IP6K2 led to stronger inhibition in *PIK3CA*-mutant cells than in wildtype counterparts (Fig. 2h). In skin cutaneous melanoma (SKCM), KO of ATP6AP1 showed preferential lethality in *PCLO*-mutant cells (Fig. 2h). Together, these findings delineate a wide array of cancer-type specific mutation-associated gene dependencies, beyond the conventional matched marker-target associations, providing opportunities for the development of novel targeted therapies.

### Profiling mutation-associated functional vulnerabilities

Building on the mutation-target dependency map, we observed that individual mutation markers were frequently associated with multiple targets across cancer types (Fig. 3a). We, therefore, hypothesized that these targets might be functionally related and involved in shared biological processes. To test this, we performed Kyoto Encyclopedia of Genes and Genomes (KEGG) pathway enrichment analysis on the sets of target genes linked to each of the mutation markers, aiming to uncover pathway-level vulnerabilities in each cancer type^27^. This analysis revealed 1-237 significantly enriched pathways (FDR≤0.05) associated with 1-68 mutation markers across 20 cancer types, which were classified into five major categories: metabolism, cellular processes, genetic information processing, environmental information processing, and organismal systems (Fig. 3b, Extended Data Figs. 5a & 6a). Focusing on pathways associated with the most frequently mutated marker in each cancer type (Extended Data Fig. 3b), a total of 70 significant enrichments were identified in 14 out of 20 cancer types such cytosolic DNA-sensing pathway in *PIK3CA*-mutant BRCA and nucleotide excision repair in *TP53*-mutant OV (Fig. 3c, Extended Data Fig. 6b). These biological processes represent high-priority and broadly applicable vulnerabilities. In contrast, no enriched pathways were found for four frequent mutation markers, namely *APC* in COAD, *TP53* in ESCA, LUSC, and READ, *PTEN* in GBM, and *BRAF* in SKCM (Fig. 3c), indicating that dominant mutations do not always guide shared functional dependencies.

**Fig. 3.**
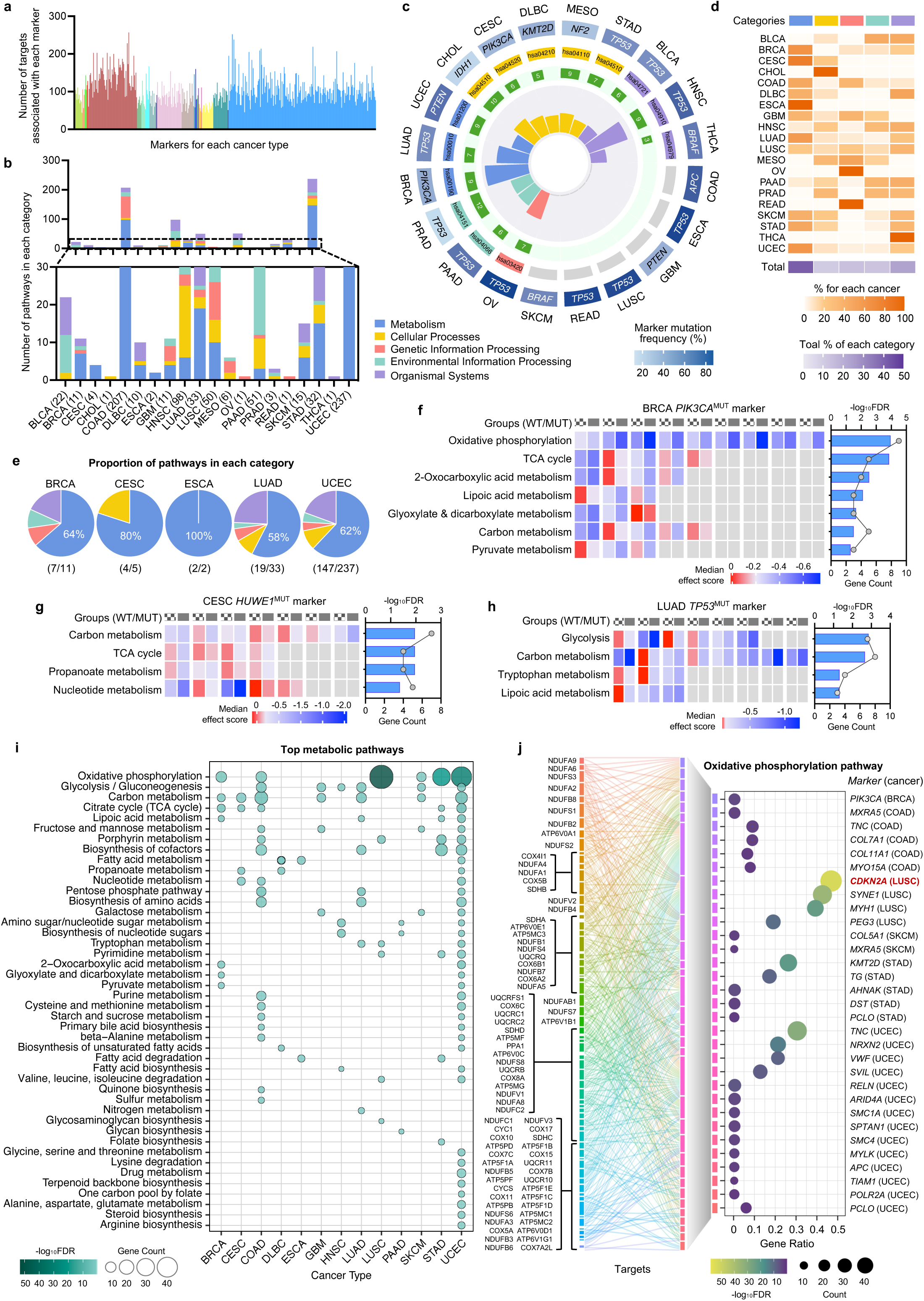

Among the five KEGG pathway categories, metabolism-related pathways emerged as the most predominant, being commonly enriched across multiple cancer types (Fig. 3d). In particular, they accounted for over 50% of all enriched pathways in BRCA, CESC, ESCA, LUAD, and UCEC (Fig. 3e). For example, targets associated with *PIK3CA* mutations, the most frequently mutated marker in BRCA, were significantly enriched in oxidative phosphorylation (OXPHOS), citrate cycle (TCA cycle), 2-oxocarboxylic acid metabolism, lipoic acid metabolism, glyoxylate and dicarboxylate metabolism, carbon metabolism, and pyruvate metabolism, with OXPHOS showing the strongest enrichment, suggesting increased bioenergetic and biosynthetic demands in *PIK3CA*-mutant BRCA cells (Fig. 3f). Similarly, in CESC, *HUWE1* mutation-associated targets were enriched in carbon metabolism, TCA cycle, propanoate metabolism and nucleotide metabolism (Fig. 3g), while glycolysis, carbon metabolism, tryptophan metabolism and lipoic acid metabolism pathways were enriched in *TP53*-mutant LUAD cells (Fig. 3h). Therefore, these results suggest that metabolic dependency is a dominant feature of mutation-defined cancer subgroups. To further delineate these associations, we examined significantly enriched metabolism pathways within each cancer type (Fig. 3i). Among the top-ranked pathways, OXPHOS emerged as the most prominent, being enriched across six cancer types (Fig. 3i).

We next investigated marker-OXPHOS associations across the six cancer types and identified 31 significant associations, with the strongest enrichment observed in *CDKN2A*-mutant LUSC (Fig. 3j). The tumour suppressor gene *CKDN2A* is frequently inactivated in LUSC, with loss-of-function mutations reported in up to 72% of cases^28^. While *CDKN2A* loss has been shown to regulate lipid metabolism in GBM, its relationship with OXPHOS remains unclear^29^. To investigate this, we assessed OXPHOS dependence in a panel of LUSC cell lines with either wildtype or loss-of-function mutations of *CDKN2A*, using both gene silencing and pharmacologic inhibition approaches. For gene silencing, we selected three representative OXPHOS targets, including UQCRB, a component of mitochondrial complex III, and NDUFS7 and NDUFS8, components of complex I, and performed siRNA-mediated gene knockdown (Fig. 4a). In line with our analysis, functional validation confirmed that *CDKN2A*-mutant cells (NCI-H520, NCI-H1703, and LK-2) exhibited marked growth inhibition upon silencing of these targets, whereas wild-type cells (NCI-EBC-1, SK-MES-1, and NCI-H2170) were minimally affected (Fig. 4b, Extended Data Fig. 7a). Consistently, pharmacologic inhibition of OXPHOS using IM156, a mitochondrial complex I inhibitor, and metformin, a first-line antidiabetic drug known to target complex I and ATP synthase, suppressed mutant cell growth to a greater extent than that of wild-type cells (Fig. 4c, d), phenocopying the effects of silencing OXPHOS targets (Fig. 4b)^30–32^. Furthermore, knockdown of CDKN2A re-sensitized wild-type cells to both IM156 and metformin treatment (Fig. 4e), suggesting that this selective vulnerability to OXPHOS inhibition is dependent on *CDKN2A* loss.

**Fig. 4.**
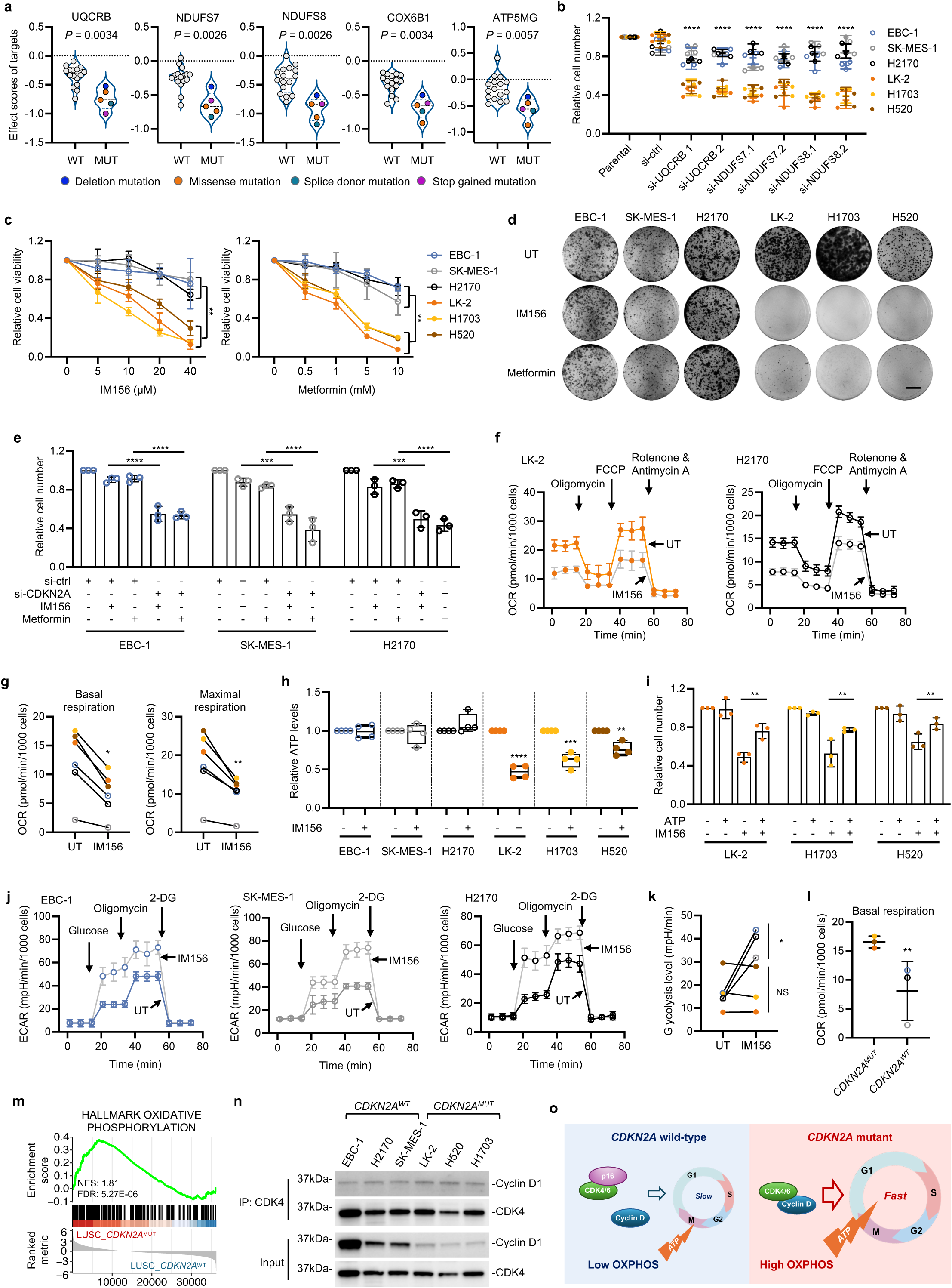

Strikingly, while treatment with OXPHOS inhibitor significantly reduced the oxygen consumption rate (OCR) in both *CDKN2A*-mutant and wild-type cell lines (Fig. 4f, g, Extended Data Fig. 7b), only mutant lines exhibited a reduction in intracellular ATP levels (Fig. 4h), and exogenous ATP supplementation at least partially rescued their impaired cell growth (Fig. 4i). In contrast, extracellular acidification rate (ECAR) measurements showed a marked increase in glycolysis in wild-type but not mutant cell lines that suggest a shift to glycolysis upon OXPHOS inhibition (Fig. 4j, k, Extended Data Fig. 7c). Together, these results suggest that *CDKN2A*-wild-type cells adapt to OXPHOS inhibition via a metabolic switch towards glycolysis to maintain energy production, whereas mutant cells exhibit limited metabolic plasticity and a heightened dependence on OXPHOS.

We next examined OXPHOS activity in both *CDKN2A*-mutant and wild-type subgroups. *CDKN2A*-mutant cell lines exhibited significantly higher basal OCR compared to their wild-type counterparts (Fig. 4l). Supportively, GSEA of TCGA LUSC RNA-seq data revealed that OXPHOS pathway was enriched in *CDKN2A*-mutant LUSC tumours compared to wild-type cases (Fig. 4m), confirming elevated OXPHOS activity in the mutant subgroup. Given that *CDKN2A* encodes the tumour suppressor p16, which arrests the cell cycle at G1 phase by sequestering CDK4/6 from binding to cyclin D, we hypothesized that *CDKN2A* mutations may lead to elevated energy demands due to unchecked cell cycle progression^33^. Indeed, co-immunoprecipitation assays showed an elevated association of CDK4 and cyclin D1 in *CDKN2A*-mutant compared to wild-type LUSC cell lines, relative to their input expression levels (Fig. 4n), supporting a model in which mutant *CDKN2A* drives OXPHOS upregulation to meet the bioenergetic demands caused by high cyclin D-CDK complexes-induced cell proliferation (Fig. 4o). Collectively, these findings establish OXPHOS as a metabolic dependency in *CDKN2A*-mutant LUSC and highlight the potential of repurposing metformin, a first-line antidiabetic medicine, to treat this mutation-defined subgroup.

### Prioritizing actionable dependencies by protein overexpression

Having identified mutation-associated targets, we next sought to evaluate their therapeutic potential by comparing protein expression levels between mutant tumours and normal tissues using CPTAC proteomic data. Among six cancer types with available mutation and proteomic data in both tumour and normal samples, we prioritized 1,080 targets that were significantly overexpressed in mutant tumours (Fig. 5a, b), suggesting these proteins are both functionally essential and therapeutically actionable. We then assessed the extent to which these overexpressed targets were shared across cancer types. Strikingly, no targets were consistently increased in protein abundance in more than four cancer types, reinforcing a largely cancer type-specific expression landscape. Nevertheless, nine targets were recurrently increased across four cancer types (Fig. 5c) and these proteins represent pan-cancer candidates with therapeutic potential. To determine whether the observed protein-level increase was transcriptionally driven, we analyzed matched RNA-seq data from CPTAC. Across four cancer types with available transcriptomic profiles, protein and RNA differential expression levels were strongly correlated overall (r = 0.6841) (Fig. 5d). Individually, HNSC, LUAD, and LUSC showed similar strong correlations (r = 0.6606, 0.6813, and 0.7170, respectively), whereas PAAD displayed a weaker correlation (r = 0.1945), likely due to fewer identified targets (Fig. 5d). Notably, over 70% of protein-overexpressed targets in HNSC, LUAD, and LUSC also exhibited consistent upregulation at the transcript level (Fig. 5e), suggesting that overexpression of these targets is predominantly driven by transcriptional activation.

**Fig. 5.**
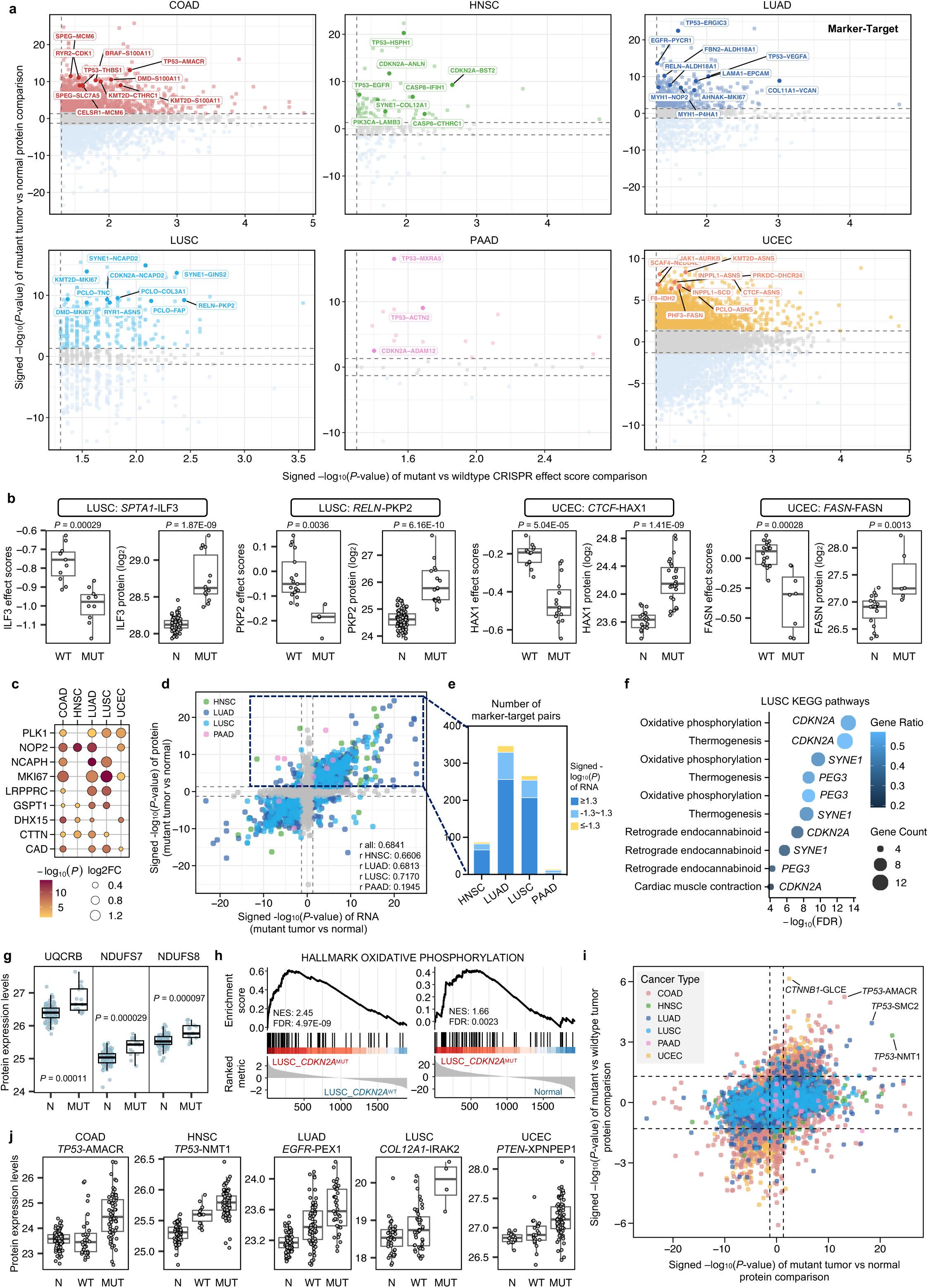

To further investigate the proteomic basis of mutation-associated vulnerabilities in LUSC, we performed KEGG pathway enrichment analysis on the sets of overexpressed targets associated with each of the mutation markers in LUSC (Fig. 5f). Remarkably, the most significant mutation-pathway association identified was *CDKN2A* mutation-OXPHOS (Fig. 5f), consistent with our earlier findings that *CDKN2A*-mutant LUSC cells exhibit selective OXPHOS dependency (Fig. 4b-d). Notably, UQCRB, NDUFS7, and NDUFS8, three OXPHOS targets validated as essential in *CDKN2A*-mutant LUSC cells (Fig. 4b), were also significantly increased at the protein level (Fig. 5g). GSEA of matched CPTAC RNA-seq data further supported these findings, showing OXPHOS pathway enrichment in *CDKN2A*-mutant LUSC tumours compared to both wild-type counterparts and normal lung tissues (Fig. 5h), echoing our earlier observations from TCGA RNA-seq analysis (Fig. 4m). Together, these findings establish that OXPHOS activation underpins a metabolic vulnerability with a therapeutic window in *CDKN2A*-mutant LUSC, highlighting that OXPHOS inhibition represents a promising strategy for treating this molecularly defined subgroup.

We next conducted comparative proteomic analysis of the identified targets between mutant and wild-type tumours to explore whether mutation-associated dependencies are driven by changes in protein abundance. Among all mutation-target associations across the six cancer types, 7-507 targets exhibited significantly higher expression in mutant tumours compared to their wild-type counterparts (Fig. 5i). Notably, 76% of these targets were also increased in mutant tumours relative to normal tissues (Fig. 5i), offering potential therapeutic windows for targeting these candidates. For example, *TP53* mutations, one of the most frequently altered mutation markers in COAD, were associated with increased protein levels of AMACR, an essential enzyme involved in the oxidation of bile acid intermediates and branched chain fatty acids (Fig. 5j)^34^. In LUAD, *EGFR*-mutant tumours showed elevated expression of PEX1, a key regulator of peroxisome assembly and function, relative to both wild-type tumours and normal lung tissues (Fig. 5j)^35^. In UCEC, XPNPEP1 was selectively overexpressed in *PTEN*-mutant tumours (Fig. 5j). Together, these analyses prioritize proteins that were overexpressed in marker-mutant subgroup, representing potential context-specific therapeutic targets for drug development.

### Linking pharmacologic sensitivities to mutation-associated dependencies

Building on our marker-target-cancer associations from CRISPR-Cas9 screening (Fig. 2g), we next extended our analysis from genetic vulnerabilities to therapeutic responses by profiling drug sensitivities using large-scale pharmacologic data from the PRISM Repurposing dataset^36^. Consistent with our previous approach, cancer cell lines were stratified into mutant or wild-type subgroups based on defined mutation markers for each cancer type (Extended Data Fig. 3b). Drug responses were then compared between the two groups across a broad panel of compounds, including 90 chemotherapies, 583 targeted drugs and 769 non-oncology agents. This analysis uncovered 4,695 mutation-drug-cancer associations across 14 cancer types, in which mutant cell lines exhibited significantly greater sensitivity to the corresponding compounds compared to wild-type counterparts (Fig. 6a). The most significant associations included enhanced sensitivity of *KMT2D*-mutant COAD cells to digitoxigenin, a cardenolide and the aglycon of digitoxin, *RB1*-mutant bladder urothelial carcinoma (BLCA) cells to a DNA topoisomerase I inhibitor 10-HCPT, and *TP53*-mutant breast invasive carcinoma (BRCA) cells to Selinexor, a selective inhibitor of nuclear export that blocks exportin 1 (Fig. 6a, b)^37,38^. To determine whether our predicted mutation-target-cancer associations are supported by drug response data, we cross-referenced compounds from the PRISM Repurposing dataset known to inhibit the targets identified from CRISPR-Cas9 screens. Although matched compounds were available for only 5.8% of associations, we identified 113 associations for which at least one matched compound exhibited significantly stronger growth inhibition in marker-mutant than in wild-type cell lines (Fig. 6c), recapitulating the effects observed in CRISPR-Cas9 screens. Therefore, these associations represent mutation-defined, actionable vulnerabilities supported by both genetic and pharmacologic evidence.

**Fig. 6.**
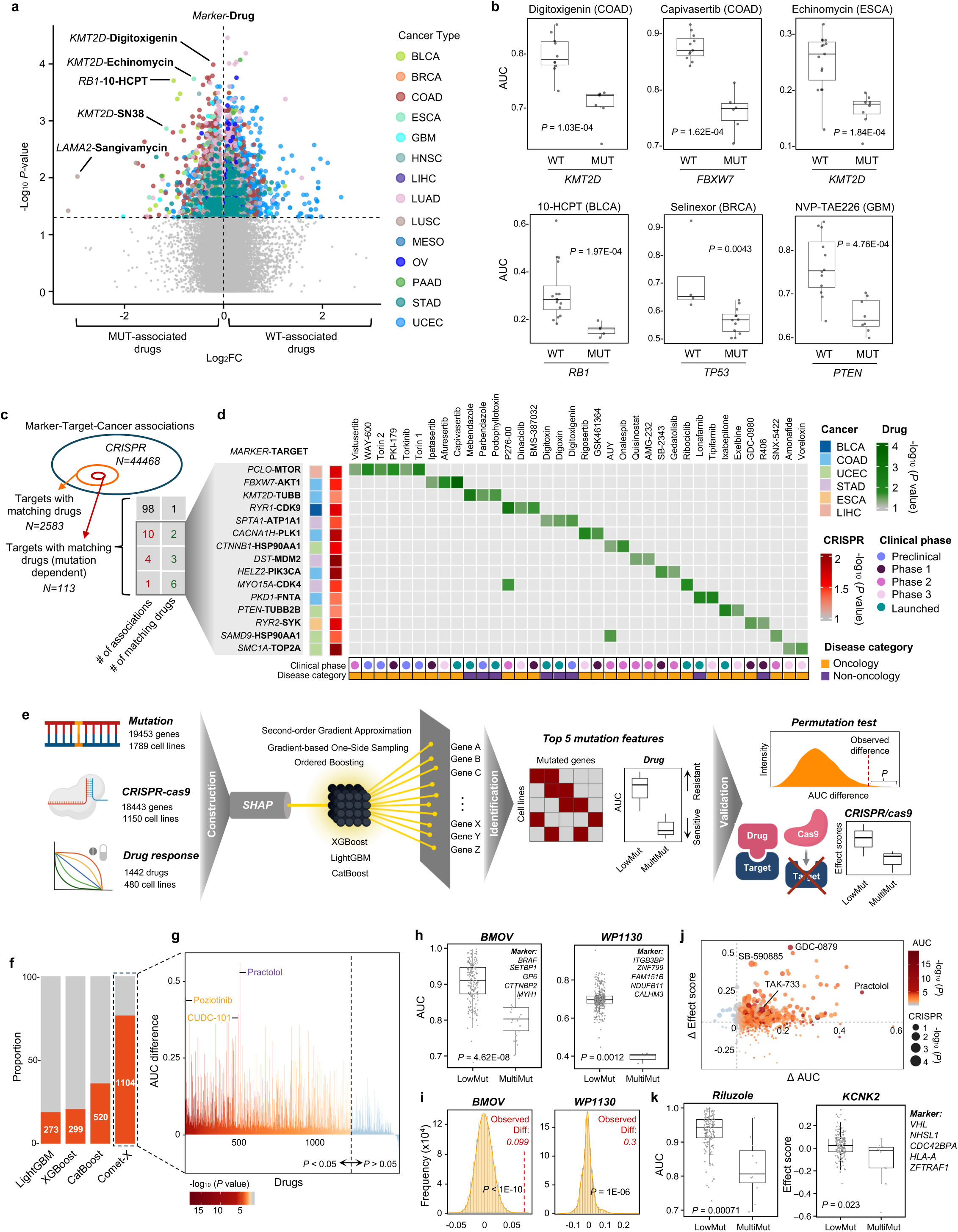

While most mutation-target-cancer associations were matched to a single compound, 15 associations had two or more compounds inhibiting the same target (Fig. 6c, d). Strikingly, in each of these associations, all matched compounds consistently demonstrated greater inhibitory effects in marker-mutant cell lines (Fig. 6d). Across these 15 associations, a total of 36 compounds were identified, of which 30 are either clinically approved or currently in clinical development (Fig. 6d). While the majority of these compounds are chemotherapeutics or targeted oncology agents, such as mTOR inhibitors Vistusertib, WAY-600, Torin 1, Torin 2, PIK-179, and Torkinib, which showed enhanced activity in *PCLO*-mutant LIHC cells, or the DNA-interacting agents Amonafide and Voreloxin, more potent in *SMC1A*-mutant UCEC, a notable subset were originally developed for non-oncology indications (Fig. 6d)^39–41^. Digitoxin and digoxin, two cardiac glycosides clinically used to treat congestive heart failure, along with their aglycone derivative digitoxigenin, showed significantly stronger inhibitory effects in *SPTA1*-mutant STAD cells than their wildtype counterpart (Fig. 6d)^42^. Moreover, the anthelmintic drugs mebendazole and parbendazole demonstrated greater inhibition in *KMT2D*-mutant COAD cells (Fig. 6d). In addition, Lonafarnib, an FDA-approved drug for Hutchinson–Gilford progeria syndrome, also showed selective activity in *PKD1*-mutant COAD cells (Fig. 6d)^43^. Taken together, these findings highlight non-oncology drugs with repurposing potential for mutation-defined cancer subgroups, offering novel therapeutic opportunities.

Given that cancer cells frequently harbor mutations in multiple genes that may interact to shape therapeutic responses, we next investigated whether combinations of mutant genes could serve as informative markers to predict drug sensitivities and target dependencies^44^. To identify combinatorial marker-drug associations that could not be predicted by single gene mutation alone, we developed a machine learning-based computational framework called Combinatorial Mutation-Enabled Therapy Exploration (Comet-X) (Fig. 6e). Comet-X integrated three independent gradient boosting models, XGBoost, LightGBM, and CatBoost, designed to uncover drug responses and CRISPR-Cas9 gene essentialities strongly associated with different combinatorial mutation patterns across pan-cancer cell lines (Fig. 6e)^45^. For each compound, Comet-X computes and integrates model-specific importance scores and SHAP values, nominating the top five predictive mutant genes (Fig. 6e). Based on these prioritized markers, 1,104 associations were identified in which cancer cell lines harbouring at least two of the top five mutant genes (MultiMut group) exhibited significantly greater drug sensitivity than those with none or only one gene mutation (LowMut group) (Fig. 6f, g, Extended Data Fig. 8a). The strongest associations included increased sensitivity to BMOV (Bis(maltolato)oxovanadium), an insulin mimetic with anti-diabetic effect, in cells with combinatorial mutations in *BRAF*, *SETBP1*, *GP6*, *CTTNBP2*, and *MYH1* (Fig. 6h), and to WP1130, a selective deubiquitinase inhibitor, in cells with mutations in *ITGB3BP*, *ZNF799*, *FAM151B*, *NDUFB11*, and *CALHM3* (Fig. 6h). Notably, the integrative strategy of Comet-X substantially outperformed individual model, yielding a greater number of significant associations across the compound panel: XGBoost, LightGBM, and CatBoost identified only 273, 299, and 520 associations, respectively, compared to 1,104 by Comet-X (Fig. 6f).

To evaluate whether the observed drug sensitivities linked to combinatorial mutations could occur by chance, we performed permutation testing with 1,000,000 random label shuffles to each of the 1104 associations identified by our framework. Remarkably, 99.3% (n = 1096/1104) of associations had observed ΔAUC values located in the extreme lower tail of their respective null distributions (permutation P < 0.05), with 53.8% (n = 594/1104), including BMOV and WP1130, reaching P < 0.001, (Fig. 6i), confirming the robustness and biological relevance of these predictions. Furthermore, among associations involving compounds with known CRISPR-Cas9-profiled targets, Comet-X identified 109 associations in which both drug sensitivity and gene knockout effects were significantly stronger in the MultiMut group compared to the LowMut group (Fig. 6j, k, Extended Data Fig. 8b). These associations encompassed 65 oncology drugs such as TAK-733, a MEK inhibitor, as well as 44 compounds originally developed for non-oncology indications (Fig. 6j, k, Extended Data Fig. 8b)^46^. For example, Riluzole, a neuroprotective drug targeting TREK-1 encoded by the *KCNK2* gene, also exhibited enhanced activity in cells carrying combinatorial mutations in *VHL*, *NHSL1*, *CDC42BPA*, *HLA-A*, *and ZFTRAF1*, recapitulating the effect of *KCNK2* KO (Fig. 6k)^47^. Together, these results demonstrate the extended value of integrative machine learning approaches in discovering combinatorial mutation markers for selective cancer vulnerabilities, extending beyond the scope of single-gene mutation markers.

## Discussion

In this study, we present a comprehensive pan-cancer framework that systematically connects hallmark gene mutations with cancer vulnerabilities by integrating multi-omic, functional, and pharmacologic datasets across diverse cancer types. Our analysis systematically links hallmark gene mutations to functional dependencies and drug responses, revealing context-specific vulnerabilities that are shaped by both single-gene and combinatorial mutation patterns. Building on this foundation, we further identify essential biological programs within genetically defined subgroups and prioritize protein targets that are both functionally essential and selectively overexpressed. Notably, our analysis highlights opportunities to repurpose non-oncology drugs for genetically stratified cancers, thereby broadening the therapeutic potential of approved drugs.

Our analysis revealed distinct, lineage-specific patterns of hallmark gene expression across cancer types. Hallmark pathway enrichment profiles similarly varied across different cancer types. These observations reinforce the concept that the activation of cancer hallmarks is not uniform across malignancies but shaped by tumour lineage, highlighting the importance of delineating hallmark vulnerabilities within individual cancer types. Importantly, hallmark gene expression profiles in cancer cell lines were largely aligned with those in tumours of matched cancer types, supporting their use as experimental models for patient tumours in probing hallmark vulnerabilities. Together, these findings establish a foundation for leveraging gene dependency and drug sensitivity data from cancer cell lines to dissect hallmark vulnerabilities within individual cancer types. Cohort-level CRISPR screens revealed both universal and cancer type-specific hallmark gene dependencies, yet substantial heterogeneity within tumour types underscored the limitations of population-level analyses. By stratifying cell lines based on recurrent, cohort-specific mutations, we achieved finer-resolution dissection of functional dependencies. This approach not only recapitulated canonical associations such as MDM2 in *TP53*-wildtype cells and PIK3CA in *PIK3CA*-mutant contexts, but also uncovered novel, functionally important dependencies exemplified by BRCA2 dependency in *DYNC1H1*-mutant colorectal cancer and HAX1 dependency in *CTCF*-mutant UCEC. These findings underscore the value of mutation-stratified analyses in uncovering hidden vulnerabilities and broadening the repertoire of therapeutic targets beyond canonical oncogenic drivers. As such, our marker-target dependency map offers a foundation to validate novel targets and advance the development of more effective, precision-guided cancer therapies.

This next step of target validation is particularly important as not all hallmark gene mutations yield functional dependencies, and some observed associations may only reflect correlation rather than causation. Mechanistic validation will be essential to define the biological basis of these links and guide therapeutic prioritization. Our approach, which stratified cell lines by the presence of mutations, did not resolve functional heterogeneity among individual mutations, such as loss-of-function and gain-of-function events, because the biological consequences of many specific hallmark gene mutations remain undefined. Future efforts incorporating functional mutation annotations will be critical to refine dependency predictions and enhance biological interpretability. Finally, because our dependency map is derived from cancer cell lines, which, while largely representative of their tumour counterparts, do not fully capture the influence of the tumour microenvironment or immune context. Thus, validations in patient-derived organoids, xenografts, or *in vivo* functional screens will be essential to translate the findings into actionable therapeutic strategies.

Building on the mutation-target dependency map, we further uncovered vulnerabilities in biological processes linked to specific mutations in different cancer types. A particularly striking finding was the selective dependency on OXPHOS in *CDKN2A*-mutant LUSC. While *CDKN2A* loss is among the most frequent alterations in LUSC, its therapeutic implications have remained elusive^28^. Our integrative analyses, supported by functional validation, demonstrated that *CDKN2A*-mutant LUSC cells exhibited heightened reliance on OXPHOS, rendering them selectively sensitive to OXPHOS inhibition. Notably, this vulnerability can be exploited by metformin, an FDA-approved antidiabetic drug, offering an immediately actionable opportunity for therapeutic repurposing.

To refine the therapeutic relevance of marker-target dependencies we discovered, we prioritized targets that were both essential in mutant tumours and significantly increased at the protein level relative to normal tissues. This dual criterion highlights expression-defined therapeutic windows and increases the tractability of these targets for drug development. Yet, it is important to note that the absence of increased protein expression does not exclude therapeutic potential. Post-translational modifications (PTMs), including phosphorylation, acetylation, and methylation, often differ markedly between tumours and normal tissues, providing alternative opportunities for intervention independent of protein abundance^49^. Moreover, elevated target protein levels in mutant versus wild-type tumours may also reflect downstream effects of the mutations, offering mechanistic insight into the observed functional dependencies. Nonetheless, protein abundance captures only part of the regulatory landscape. Functional vulnerabilities may also arise from PTMs that modulate protein activity, subcellular localization, or interaction dynamics. Therefore, incorporating PTM proteomics data will be critical to uncover these additional regulatory layers and expand the spectrum of actionable targets.

By integrating drug sensitivity profiles with mutation-defined dependencies, we identified compounds, including oncology and non-oncology agents, that target the mapped vulnerabilities and showed selective activity in the corresponding mutant-defined subgroups. These findings open opportunities for repurposing non-oncology drugs for cancer treatment, as a rapid and cost-effective strategy, particularly for cancer types with limited targeted therapies^50–52^.

A key innovation of this work distinguishes it from earlier progress in integration of multi-omic and functional data, such as studies focusing on target identification driven by protein overexpression and activation^53^. Tumorigenesis and progression select for phenotypes that result from, or are marked by, complex mutation genotypes and signatures, collectively shaping oncogenic signalling and therapeutic responses. So we ensured that this analysis could exploit that genetic complexity. Our machine learning-based framework, Comet-X, was designed to search a space beyond single-gene markers to unrestricted combinations of hallmark gene mutations. It prioritizes the most relevant mutation events and uncovers therapeutic opportunities associated with combinations of mutation markers. Due to the limited number of cell lines within individual cancer types, Comet-X was necessarily performed in a pan-cancer setting, as machine learning models require sufficiently large sample size to compute reliable predictions^54^. As larger datasets become available, Comet-X will be well placed to predict further cancer type-specific associations, further enhancing its utility in precision oncology.

In conclusion, we have developed a comprehensive resource of hallmark gene marker-target associations alongside a compendium of potentially effective drugs for mutation-defined cancer subgroups. These datasets provide a robust foundation for understanding the biological impact of hallmark gene mutations, validating novel targets for precision oncology, developing new targeted therapies, and repurposing existing drugs for more effective cancer treatment.

---

**Extended Data Fig. 1.**
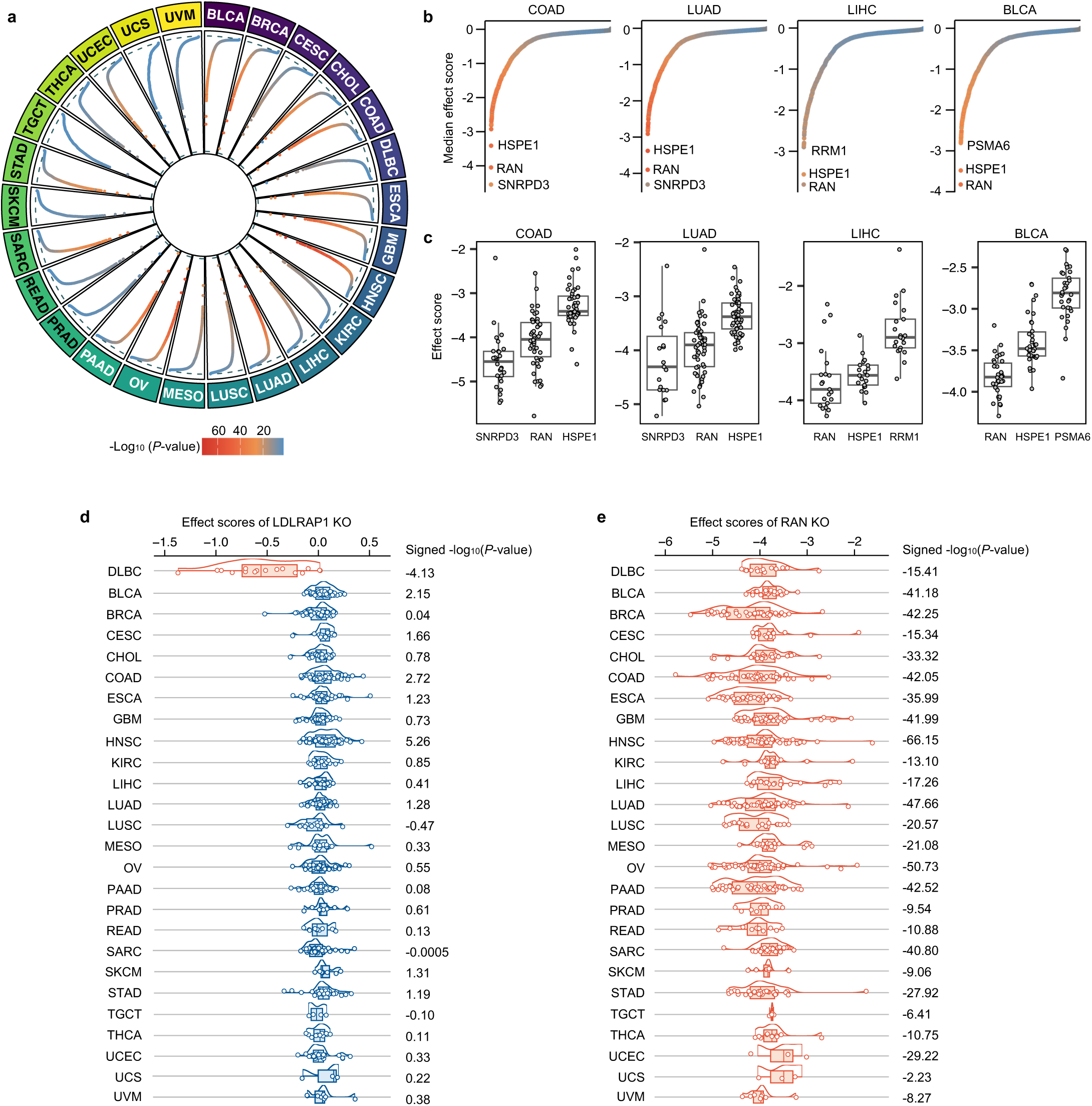

**Extended Data Fig. 2.**
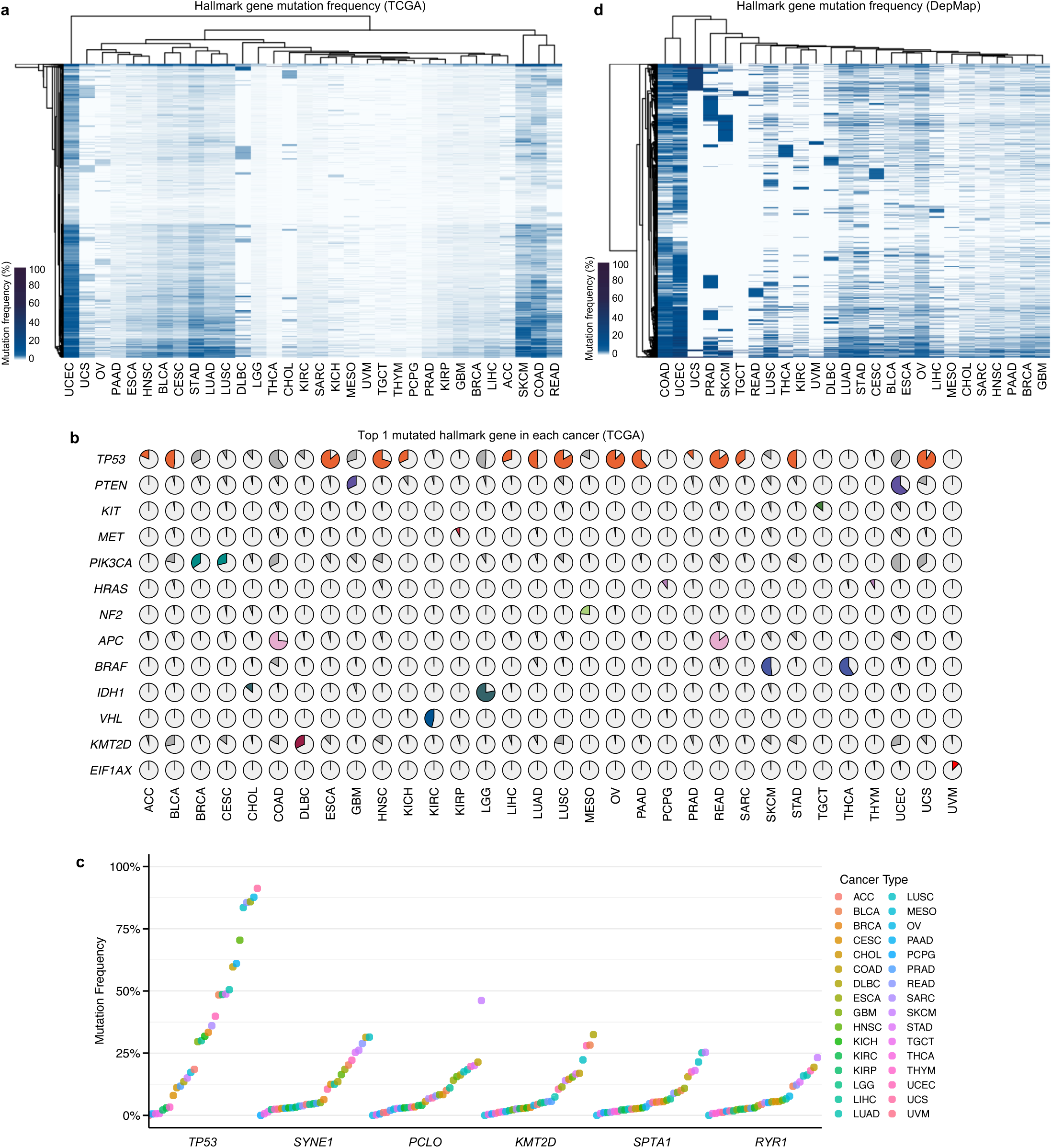

**Extended Data Fig. 3.**
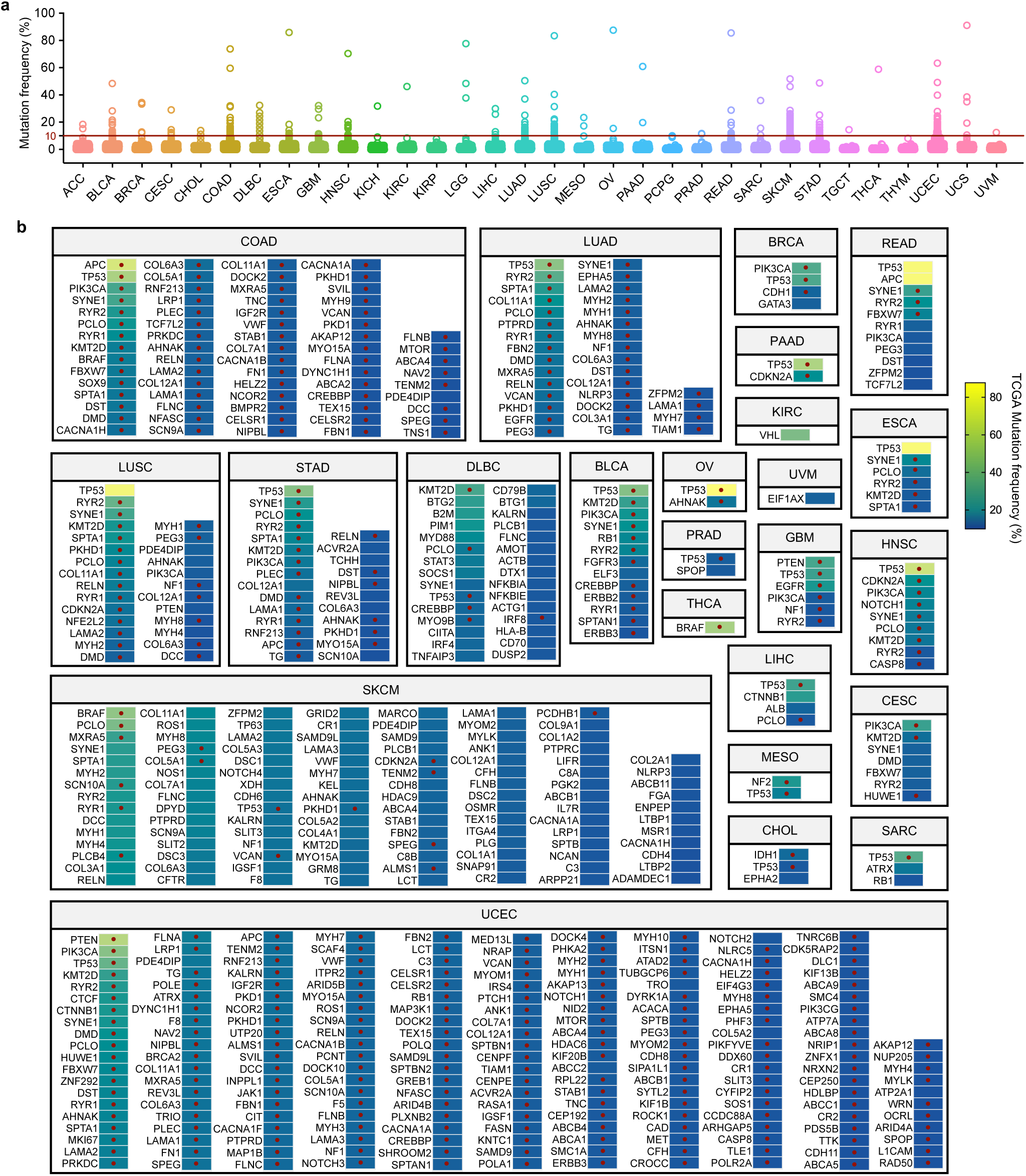

**Extended Data Fig. 4.**
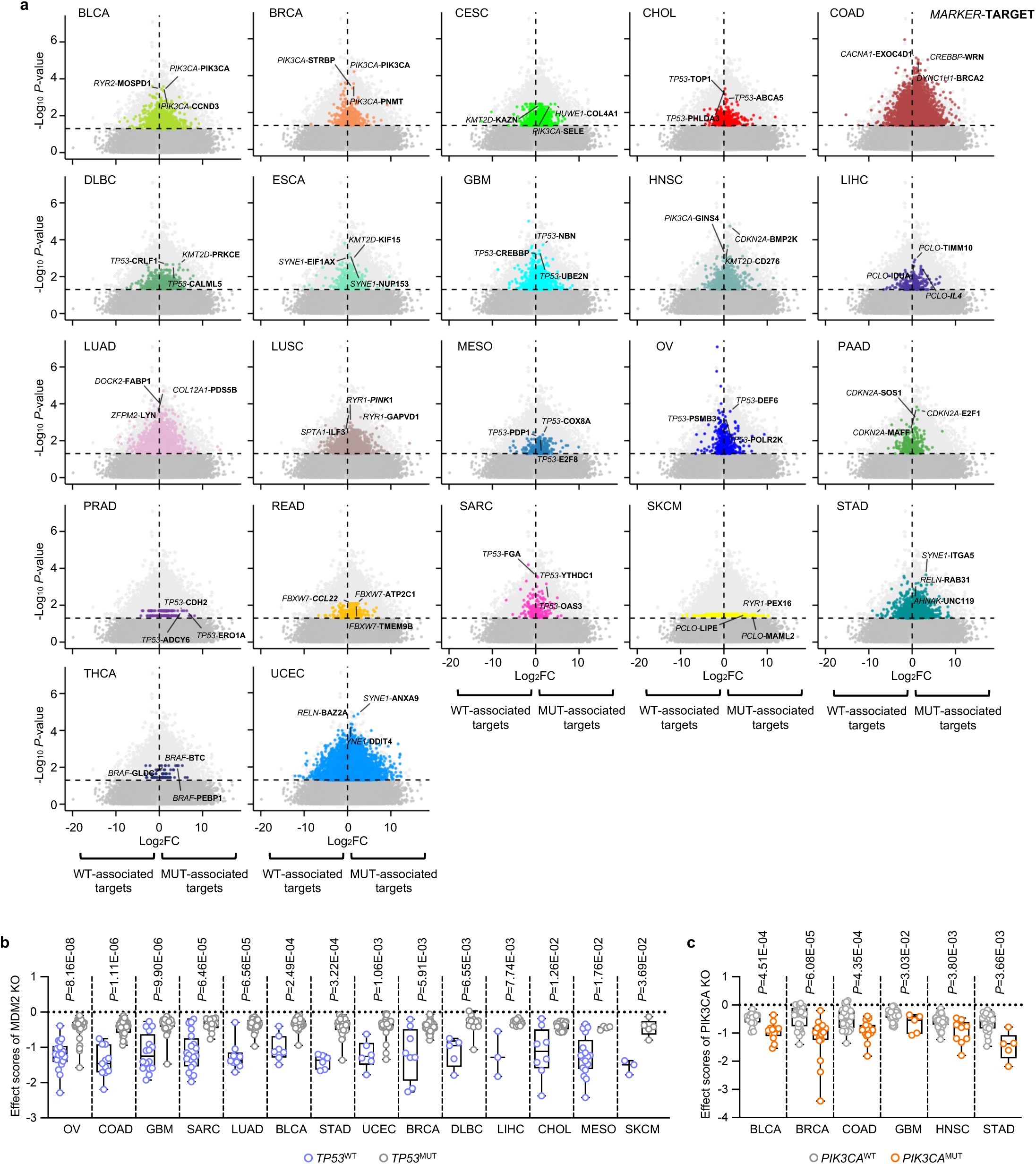

**Extended Data Fig. 5.**
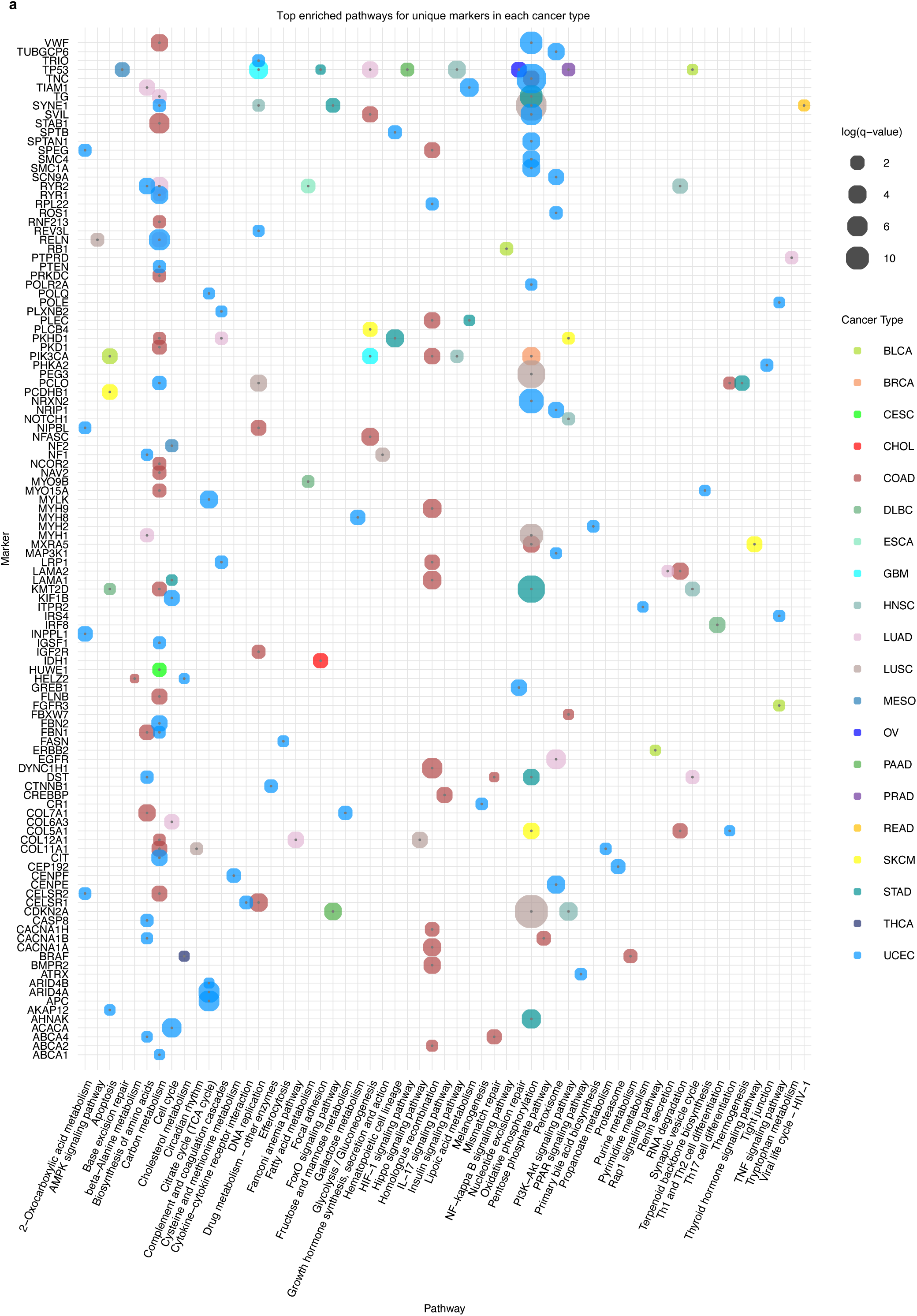

**Extended Data Fig. 6.**
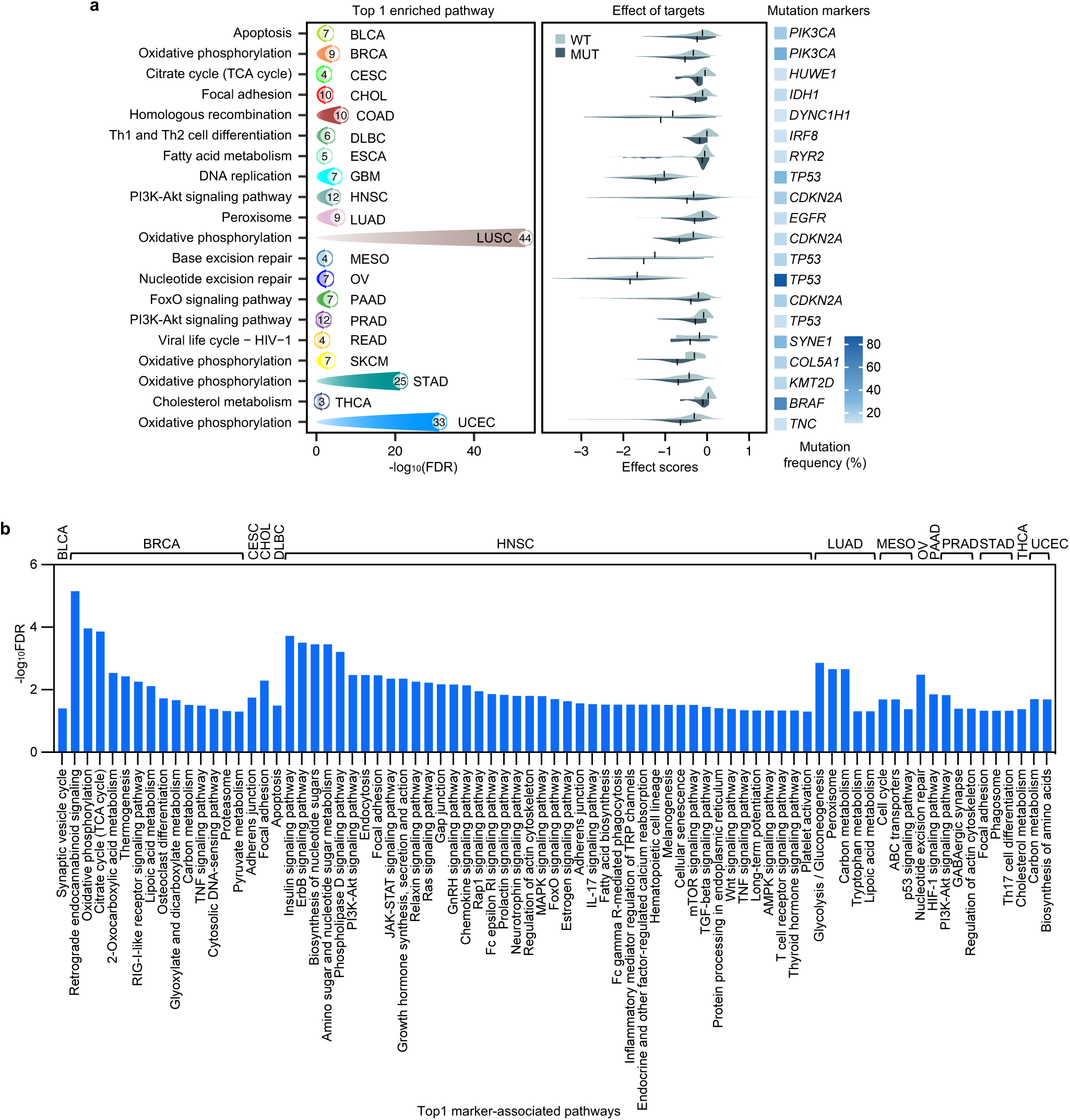

**Extended Data Fig. 7.**
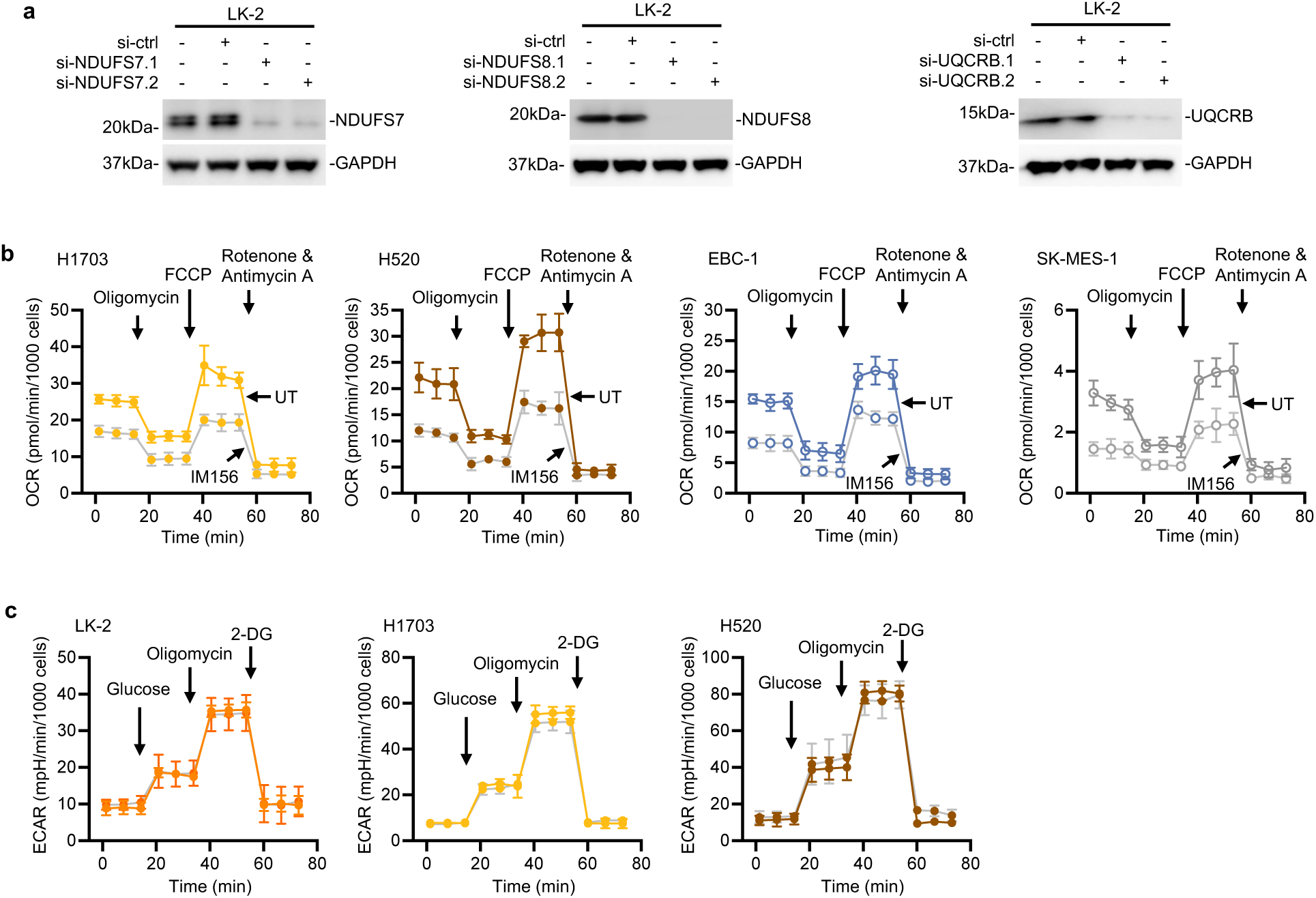

**Extended Data Fig. 8.**
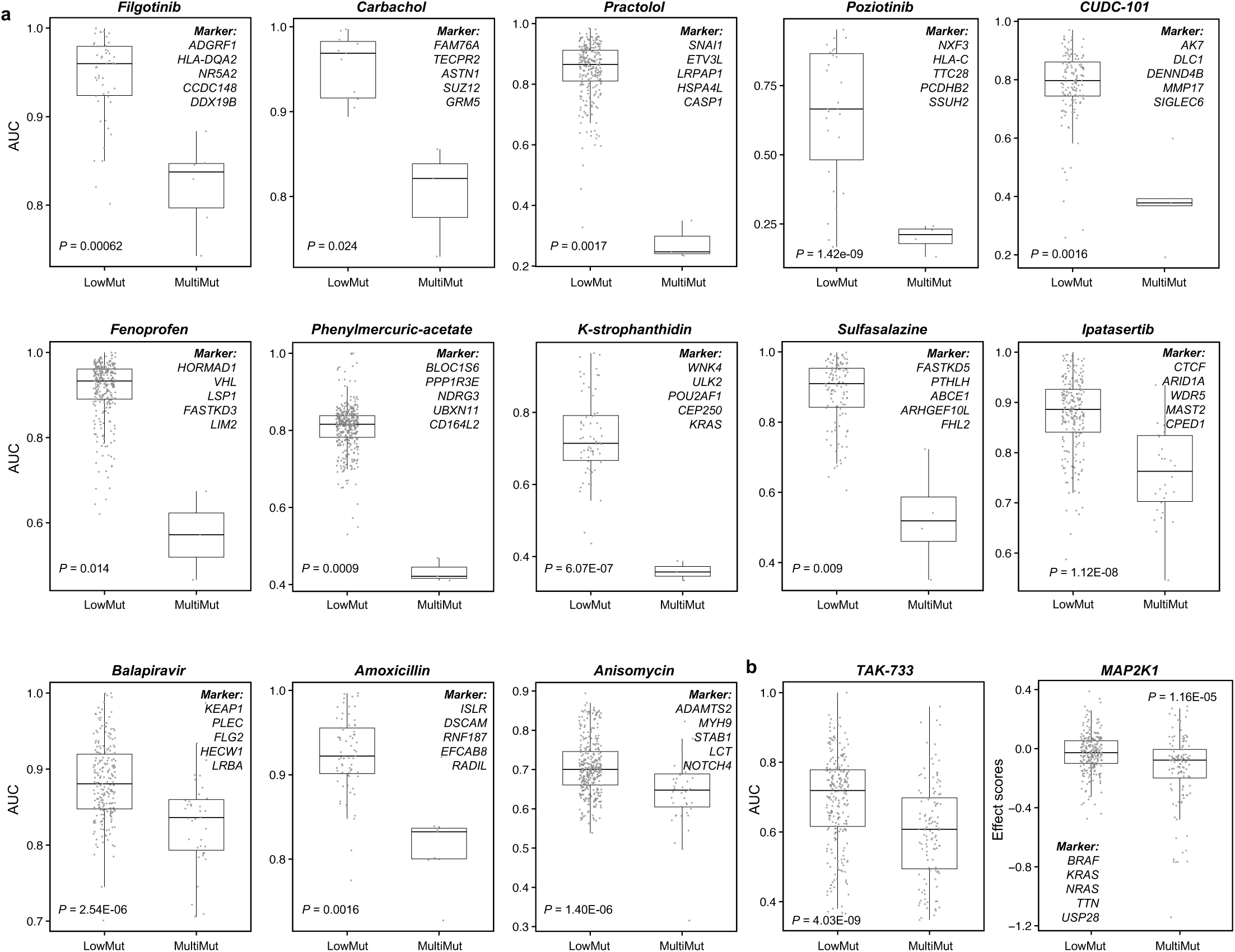

## Notes

### Competing Interest Statement

The authors have declared no competing interest.

### Funding

- National Health and Medical Research Council: 2026530

